# Stronger evidence for relaxed selection than adaptive evolution in high-elevation animal mtDNA

**DOI:** 10.1101/2024.01.20.576402

**Authors:** Erik N. K. Iverson, Abby Criswell, Justin C. Havird

**Affiliations:** Department of Integrative Biology, the University of Texas at Austin, Austin, TX, United States

## Abstract

Mitochondrial (mt) genes are the subject of many adaptive hypotheses due to the key role of mitochondria in energy production and metabolism. One widespread adaptive hypothesis is that selection imposed by life at high elevation leads to the rapid fixation of beneficial alleles in mtDNA, reflected in the increased rates of mtDNA evolution documented in many high-elevation species. However, the assumption that fast mtDNA evolution is caused by positive, rather than relaxed purifying selection has rarely been tested. Here, we calculated the *d*_N_/*d_S_* ratio, a metric of nonsynonymous substitution bias, and explicitly tested for relaxed selection in the mtDNA of over 700 species of terrestrial vertebrates, freshwater fishes, and arthropods, with information on elevation and latitudinal range limits, range sizes, and body sizes. We confirmed that mitochondrial genomes of high-elevation taxa have slightly higher *d*_N_/*d_S_* ratios compared to low-elevation relatives. High-elevation species tend to have smaller ranges, which predict higher *d*_N_/*d_S_* ratios and more relaxed selection across species and clades, while absolute elevation and latitude do not predict higher *d*_N_/*d_S_*. We also find a positive relationship between body mass and *d*_N_/*d_S_*, supporting a role for small effective population size leading to relaxed selection. We conclude that higher mt *d*_N_/*d_S_* among high-elevation species is more likely to reflect relaxed selection due to smaller ranges and reduced effective population size than adaptation to the environment. Our results highlight the importance of rigorously testing adaptive stories against non-adaptive alternative hypotheses, especially in mt genomes.

## Introduction

Altitudinal gradients are natural laboratories for ecology because they encompass overlapping clines in many environmental factors. Organisms must cope with concordant global changes in temperature, oxygen availability, and UV radiation with elevation, as well as local clines in precipitation and primary productivity. Accordingly, adaptation to high-altitude is apparent in the genetic, physiological, and ecological traits of both animals and plants (Gutierrez-Pinto et al., 2020; Projecto-Garcia et al., 2013; Read et al., 2014; Storz, 2021; Storz & Scott, 2019a; Ye et al., 2020; Zhu et al., 2018). Adaptations to hypoxia and low temperature at increased elevation have been extensively studied, particularly in birds and mammals, because of presumed selection pressure these factors exert on the high aerobic metabolism of active endotherms (Chappell et al., 2007; Cheviron & Brumfield, 2012; Gutierrez-Pinto et al., 2020; Nabi et al., 2021; Scott et al., 2015; Storz & Scott, 2019b).

Several phenotypes strongly associated with elevational adaptation, including changes in hemoglobin activity, body insulation, or behavior, are likely to be largely or entirely linked to variation in the nuclear genome. However, the process of oxidative phosphorylation, the key component of aerobic metabolism, involves the coordinated activity of genes from both the nuclear and mitochondrial genomes (Rand et al., 2004; Sloan et al., 2018; Moran et al. 2024). The electron transport system which powers oxidative phosphorylation generates both the majority of cellular ATP and a significant portion of endotherm body heat via non-shivering thermogenesis. Thus, electron transport system proteins, and mitochondrial structures and processes more generally, have emerged as key foci in studies of adaptation to elevation (Hill, 2019; Luo et al., 2013; Murray & Horscroft, 2016; Scott et al., 2018).

While some studies have proposed mechanistic adaptive hypotheses for specific substitutions in mitochondrial genes at high elevations (see Ji et al., 2012; Kostin & Lavrenchenko, 2018; Scott et al., 2011), studies more often report broad molecular signatures of selection and then assume their functional salience for high-altitude living without further tests. For instance, mitochondrial adaptation to elevation has repeatedly been inferred from unique or excess non-synonymous substitutions in mammals (Hassanin et al., 2009; Luo et al., 2008; Ning et al., 2010; Xu et al., 2005; Yu et al., 2011), birds (Zhou et al., 2014), reptiles (Jin et al., 2018), amphibians (Wang et al., 2021), freshwater fishes (Li et al., 2013), and arthropods (Li e al., 2018; Yuan et al., 2018; Zhang et al., 2013). This has often been reported via elevated ratios of non-synonymous to synonymous substitution rates—the *d*_N_/*d*_S_ ratio. While positive selection can explain excessive nonsynonymous substitutions, so can relaxed selection, a distinction which is rarely acknowledged in even the most exhaustive studies of this genre (e.g., Wang et al., 2021; see Zwonitzer & Iverson et al., 2023). Typically, the *d*_N_/*d*_S_ ratio alone cannot distinguish positive and relaxed selection from one another and additional metrics are needed (Wertheim et al., 2015).

Studies have long highlighted the sensitivity of *d*_N_/*d*_S_ to relaxed selection, particularly due to reduced effective population size (*N_e_*) (Kliman et al., 2000; Woolfit & Bromham, 2003). Relaxed selection provides an alternative explanation for substitution patterns at high-elevation because high-elevation species should often have lower *N_e_* than their low-elevation relatives. Typically, less habitat is available at higher elevations (see Eakins & Sharman, 2012), and, all else being equal, species with less habitat should have smaller ranges, lower population sizes, and experience stronger genetic drift and less efficient selection. In addition, high-elevation populations should be more fragmented by topography, which might further reduce *N_e_* (e.g, Hahn et al., 2012; Polato et al., 2017). Among possible markers, animal mitochondrial genes may be especially affected by changes in *N_e_*. In contrast to the nuclear genome, animal mtDNA lacks recombination, is uniparentally inherited, and is effectively haploid, making it more susceptible to genetic drift than nuclear loci according to classic theory (Lynch & Blanchard, 1998; Neiman & Taylor, 2009; but see Bazin et al. 2006, Edwards et al., 2021). Formal tests that can disentangle positive from relaxed selection are now available (Wertheim et al., 2015) and can be used to investigate molecular patterns of adaptation related to altitude (e.g., Gutiérrez et al., 2023; Figure 1A).

**Figure 1.**
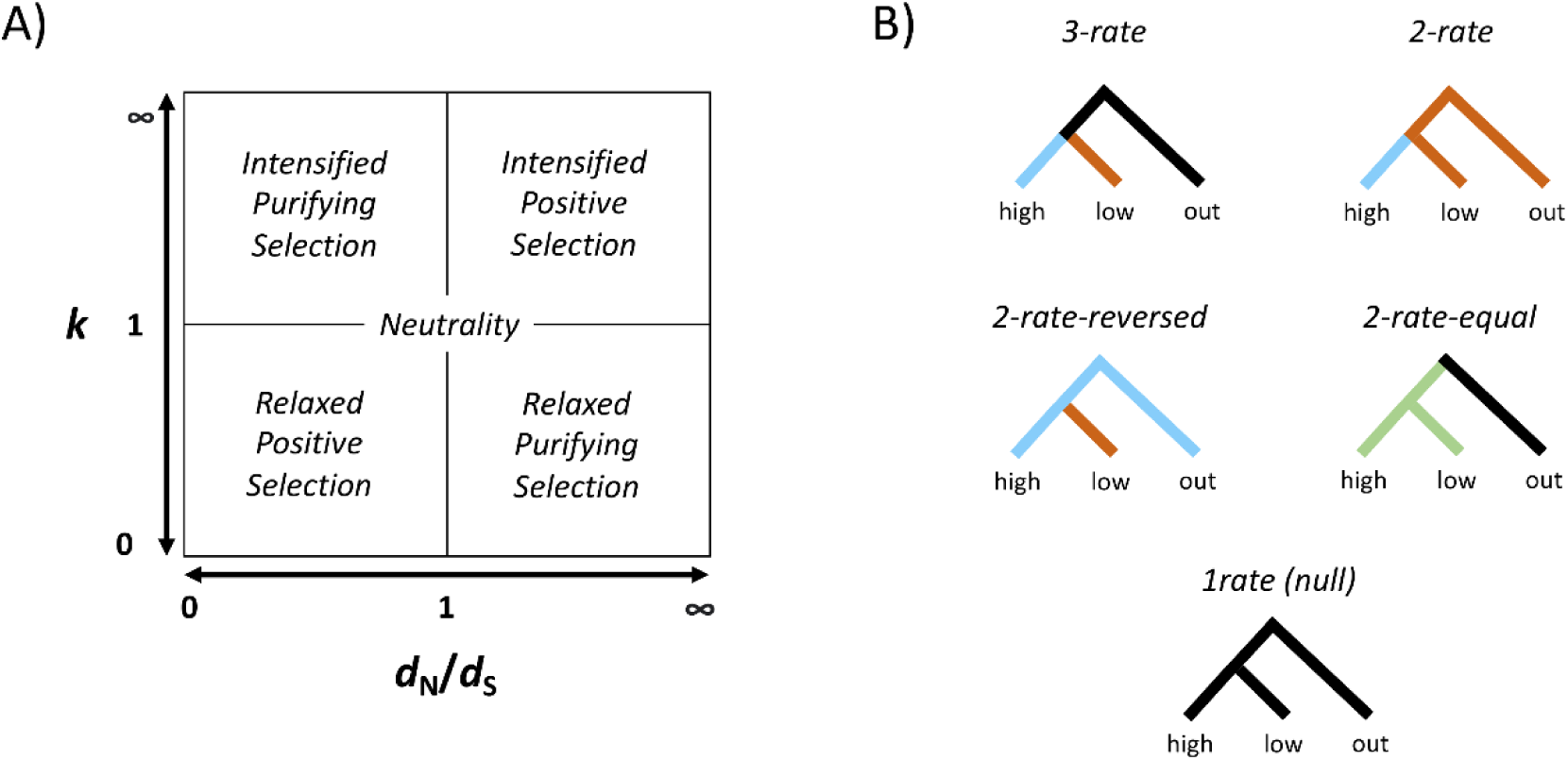
**A)** Schematic representation of selection inference from both *d*N/*d*S ratios (calculated by codeML in PAML) and *k*-values (calculated in RELAX). **B)** Illustration of different models assessed with PAML.

To foster a mechanistic understanding of adaptive and relaxed mitochondrial evolution across elevation, patterns of molecular evolution should be evaluated with regards to taxonomic group, metabolic strategy (endotherm vs. ectotherm), and body size. The impact of temperature should also be parsed from that of oxygen limitation by examining latitude as a co-variate. To that end, we examined signatures of positive and relaxed selection in all mitochondrial-encoded (mtDNA) protein-coding genes across nearly 800 species that span a range of elevations. We combined our analyses with high-quality ecological data to characterize the genetic consequences of elevation, latitude, range size, and body size across terrestrial vertebrates, freshwater fishes, and arthropods. We took two approaches; first we compared high-elevation species to their low elevation sister taxa and outgroups in 154 phylogenetically independent triads, estimating selection on high and low branches and comparing it to ecological data. We then examined 6 family or sub-order level clades totally 380 species, evaluating selection on tips at progressively higher elevations and latitudes or smaller range sizes. Together, these analyses provide multiple streams of evidence for distinguishing between adaptive evolution and neutral processes in relation to elevation and other ecological factors.

## Methods

### 1. Selection on mtDNA in focal high-elevation species

#### 1.1 Identification of high-low-outgroup triads

We first tested for selection in mtDNA of high-elevation focal species compared with their low-elevation relatives. We constructed phylogenetic trees of three-species (“triads”) consisting of a high-elevation taxon, a low-elevation sister taxon, and a low-elevation outgroup (**Supplementary Table S1**). To be included in our analyses, the high- and low-elevation taxa had to differ in maximum elevation by at least 500 m. The outgroup was the most closely-related taxon for which a mitogenome could be recovered which also had a maximum elevation 500 m or more lower than the high-elevation focal species. High- and low-elevation taxa were typically members of the same genus but sometimes were closely-related genera, subspecies, or discrete populations. The high-elevation taxon in each comparison represented what was inferred to be an independent origin of higher-elevation living compared to the other triads based on the distribution of elevation ranges among its close relatives and any available information on the history of the group. Each comparison was thus internally phylogenetically controlled and both high- and low-elevation taxa were independent of high- or low-elevation taxa in any other comparison. Care was taken to pair species that were matched for latitude, aridity, migratory behavior, and other potentially important ecological factors (**Supplemental Methods S1.1**).

#### 1.2 Analysis of selection on high-elevation species

Triads were organized into six taxonomic groups: mammals (*n* = 55 triads), birds (*n* = 34), non-avian reptiles (*n* = 13), amphibians (*n* = 23), bony fishes (*n* = 13), and arthropods (*n* = 16). Mitogenomes were downloaded from GenBank or assembled *de* novo from SRA data with MitoFinder v 1.4 (Allio et al., 2020) using default settings and the most closely-related species with a published mitogenome as a template (**Supplementary File S1**). The thirteen mitochondrial protein-coding genes for all species of a particular taxonomic group were aligned by MUSCLE (Edgar, 2004), cleaned of start and stop codons, and concatenated in alphabetical order. Then the three species of each triad were parsed into 154 individual alignments in which selection was analyzed.

Individual genes and concatenated sets of 13 genes (i.e., the mt CDS) were analyzed using codeML in PAML v. 4.7 (Yang, 2007) to calculate branch-specific values of the *d*_N_/*d*_S_ ratio, a measure of nonsynonymous substitution bias. Commonly, *d*_N_/d_S_ ratios less than one are thought to indicate purifying selection, ratios above one to indicate positive selection, and ratios close to one to represent neutrality, although this is conservative when considering an entire CDS because all mt genes are expected to be under strong purifying selection (*d*_N_/*d*_S_ < 1; **Figure 1A**). As such, our interest was mainly in relative increases or decreases in *d*_N_/*d*_S_ among related species and the relationship between relative *d*_N_/*d*_S_ and ecological variables, rather than absolute *d*_N_/*d*_S_ values. For each triad, we calculated *d*_N_/*d*_S_ on the concatenated CDS under five different models (**Figure 1B**): 1) a three-rate model giving the high and low taxa independent rates relative to the combined ancestral and outgroup branch, 2) a two-rate model grouping the low taxon with the outgroup and ancestral branch, 3) a “two-rate-reversed” model grouping the high taxon with the outgroup and ancestral branch, 4) a “two-rate-equal” model grouping the high and low taxa and their ancestral branch together, relative to the outgroup, and 5) a one-rate (null) model with the same *d*_N_/*d*_S_ applied to all branches (**Figure 1B**). We removed any triad where *d*_N_/*d*_S_ of the whole CDS on any branch was calculated to be greater than 1, an unrealistically high value for a whole mt CDS. Each triad had a minimum of 5/13 genes available for analysis in all three species. Model suitability was assessed with likelihood ratio tests and Akaike’s Information Criterion (AIC). We also used the three-rate model to calculate *d*_N_/*d*_S_ for each gene individually within each triad. We removed all runs for individual genes with predicted *d*_N_/*d*_S_ values for any species, under any model, that were either 0.001 or 999, indicating insufficient genetic variation to calculate an accurate value.

The concatenated CDS was then used as the basis for running RELAX (Wertheim et al., 2015), a phylogenetic test for relaxed selection, on the Datamonkey webserver (https://datamonkey.org/relax) . The high-elevation taxon was selected as the test branch and all other branches were reference branches. RELAX examines the distribution of *d*_N_/*d*_S_ values among sites on the test branch compared to the distribution in the reference branches. The assumption is that when selection is relaxed, *d*_N_/*d*_S_ values converge towards one, while when selection is intensified, some sites experience intensified positive selection and others experience intensified purifying selection. Thus, the distribution of *d*_N_/*d*_S_ values moves away from one and towards both extremes on test branches when intensified selection is responsible for elevated *d*_N_/*d*_S_ values. RELAX calculates a *k*-value, with *k*-values below one indicating relaxed selection on test branches, and values above one indicating intensified selection (**Figure 1A**). The program then performs a likelihood ratio test for the preferred *k*-value relative to a null model where test and reference branches have the same distribution of *d*_N_/*d*_S_ values.

#### 1.3 Analysis of ecological data

For each species in each triad in all six taxonomic groups, elevational range limits were taken from IUCN and/or several other expert sources or determined from analysis of GBIF records, range maps, and habitat information (**Table S1**). We used the highest maximum and lowest minimum reputable elevational limits for analyses and calculated midpoint elevations and latitudes, with midpoint latitude for species with ranges that straddle the equator taken as the point midway between the equator and the absolute maximum latitude. For mammals, birds, reptiles, and amphibians, IUCN shapefiles were downloaded from https://iucnredlist.org when available and latitudinal limits and range sizes were extracted using a custom Python script (see **Data Availability; File S1**). If IUCN shapefiles were unavailable, latitudinal limits and range size were estimated from available range maps, habitat distributions, and descriptive information where possible. Latitudinal limits and range size were not calculated for fishes and arthropods because shapefiles and expert range maps were typically unavailable. Body size was recorded from one of several published databases or drawn from the literature. Body size was recorded as mass for mammals and birds and log-transformed for analysis, and as snout-vent length for reptiles and amphibians and standard length for fishes. Size was not recorded for arthropods as it was typically unavailable.

#### 1.4 Statistical analyses

We compared *d*_N_/*d*_S_ between high and low branches under the three-rate model (**Figure 1B**) using paired, one-sided Wilcoxon signed-rank tests. Comparisons of the full mt CDS and individual genes were made within taxonomic groups with significance evaluated both before and after Bonferroni correction by the number of independent tests. We used Phylogenetic Generalized Least Squares (PGLS; Grafen, 1989) implemented with the R packages APE (v. 5.7.1) and phangorn (v. 2.11.1) to test for associations between *d*_N_/*d*_S_ values or *k*-values and various ecological factors. For the correlation matrix in PGLS, we used a phylogeny of the 138 high-elevation vertebrate species from each triad inferred via maximum likelihood based on a MUSCLE-generated alignment (Edgar, 2004) of amino acid sequences from all mt CDS input into RAxML v. 8 using default parameters under a PROTGAMMAWAG model (Stamatakis, 2014; **File S2**). We also used PGLS to test for relationships between both absolute and relative *d*_N_/*d*_S_ or *k* and ecological factors. In other words, for relative values the predictor variable was the ratio between high- and low-taxon values for the ecological factor, and the response was the ratio between high- and low-taxon values for *d*_N_/*d*_S_ or *k*. Ecological factors examined were absolute maximum, minimum, and midpoint elevations, maximum, minimum, and midpoint latitude, range size, and body size of the high-elevation species, as well as these values relative to values for the low-branch species. Variables were natural log-transformed when necessary to meet assumptions of normality.

### 2. Selection on mtDNA across wide-ranging clades

#### 2.1 Assembly of clades

To complement our analyses on closely-related triads and more generally examine selective pressures on mtDNA across elevation, latitude, and range size, we also surveyed 6 family- or suborder-level clades. These groups were characterized by abundant mitogenomes in NCBI and Eurasia-centered distributions typically including both the Tibetan plateau and a high-degree of latitudinal variability. Unlike in the triad-based analyses of focal high-elevation taxa, species were only included if they had complete mitogenomes in NCBI, IUCN shapefiles, and elevational data (**Supplementary Table S2**). This meant that no freshwater fish or arthropod groups were analyzed. There were three endotherm groups, Cercopithecidae (Old World monkeys; n = 72 species), Bovidae + Moschidae (sheep, goats, cattle, and musk deer; *n* = 88), and Phasianoidea (Pheasants, New World quail, guineafowl, and curassows; *n* = 67), as well as three ectotherm groups, Cryptobranchoidea (primitive salamanders; *n* = 34), Ranoidea (true frogs and relatives; *n =* 65), and Lacertoidea (wall lizards and whiptails; *n* = 55). We downloaded mitogenomes and aligned them by clade as above. We used RAxML v. 8 to construct phylogenies for each group from an alignment of the thirteen mt protein-coding genes as above, except using a GTRGAMMA model on a nucleotide alignment for the Bovidae + Moschidae (due to low amino acid divergence, last common ancestor: 16-19 Ma; Bibi, 2013) with six to seven outgroup taxa included for each group (**Table S2; File S2**).

#### 2.2 Analysis of ecological data

In addition to extracting elevational limits from the literature and extracting latitudinal limits and range size from IUCN shapefiles as in the triad analyses, we calculated a measure which we call the Climatic Index (CI), which combines latitude and elevation to express a species’ position in both climatic dimensions relative to the rest of its clade:

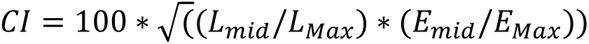

where *L_mid_* is the latitudinal midpoint of a given species’ range and *E_mid_* is its elevational midpoint, while *L_Max_* and *E_Max_* are the maximum latitude and elevation attained by *any* species in the group. Thus, the dimensionless *CI* expresses with one number how close a species’ midpoints are to the theoretical maxima for its clade (e.g, a species with elevational and latitudinal midpoints at 50% of the theoretical maximum would have a *CI* of 50.)

#### 2.3 Analysis of selection across clades

We ran codeML in PAML for each group by binning species into rate categories based on midpoint elevation, midpoint latitude, *CI*, or range size (6 groups x 4 parameters = 24 runs). Terminal branches of the phylogeny were assigned to bins of width 500 m for midpoint elevation, 5° for midpoint latitude, 5 dimensionless units for climatic index, and base-*e* exponents for range size in km^2^. These multi-rate analyses thus had between 7-13 rate bins with 1-27 species (terminal branches) per bin. All internal branches were unlabeled, making them one binned “background” rate for the analysis. We also ran codeML on a one-rate (null) model with no labeled branches, which produced a value close to that for the “background” bin of the multi-rate analysis. *d*_N_/*d*_S_ values for each bin were divided by the tree-wide *d*_N_/*d*_S_ values for the null, one-rate model and plotted as a regression over the parameter of interest. The significance of the binning scheme relative to the null model was assessed using likelihood ratio tests. We then performed RELAX on each bin of the aforementioned phylogenies, treating the binned taxa as “test” branches and the internal branches – corresponding to the unlabeled “background” *d*_N_/*d*_S_ category – as the “reference” branches, to test for relaxed selection as a function of different ecological factors.

## Results

### High-elevation taxa have elevated d_N_/d_S_ relative to low-elevation sister taxa

In total, we parsed, aligned, and concatenated 792 mt CDS including 42 novel mt CDS derived from publicly available transcriptomes, UCE, and WGS datasets (**Tables S1, S2; File S1**). We first calculated *d*_N_/*d*_S_ and *k*-values on these CDS for 154 three-species triad comparisons (**Table S3; File S3**). In 95 out of 154 triads (61.7%), the three-rate model was preferable to the one-rate model by likelihood ratio test at *P* = 0.05. In 58/95 of these triads (61.1%), the high-elevation taxon had a higher *d*_N_/*d*_S_ value. After Bonferroni correction (*P* = 0.003) the three-rate model was preferred in 40 out of 154 triads (26.0%), with 21/40 (52.5%) having higher *d*_N_/*d*_S_ in the high-taxon. *d*_N_/*d*_S_ values for concatenated alignments were always less than 1.0, indicating purifying selection in line with expectations for animal mitochondrial genomes as a whole (**Fig. 2A,C, Table S4**). Individual genes showed greater variability, including values over 1.0 (**Fig. 2B, Table S4**). Across all triad comparisons, the median *d*_N_/*d*_S_ value on the high branch was 10.9% greater than that on the low branch (*P* = 0.006, *V* = 7345.8; paired, one-sided Wilcoxon signed-rank test; **Fig. 2C**). A similar trend was found in the 95 comparisons where the three-rate model was preferred over a one rate model at *P* = 0.05 (7.1% higher on the high branch; *P* = 0.019, *V* = 2837, *d.f.* = 153; **Fig 2C**).

**Figure 2.**
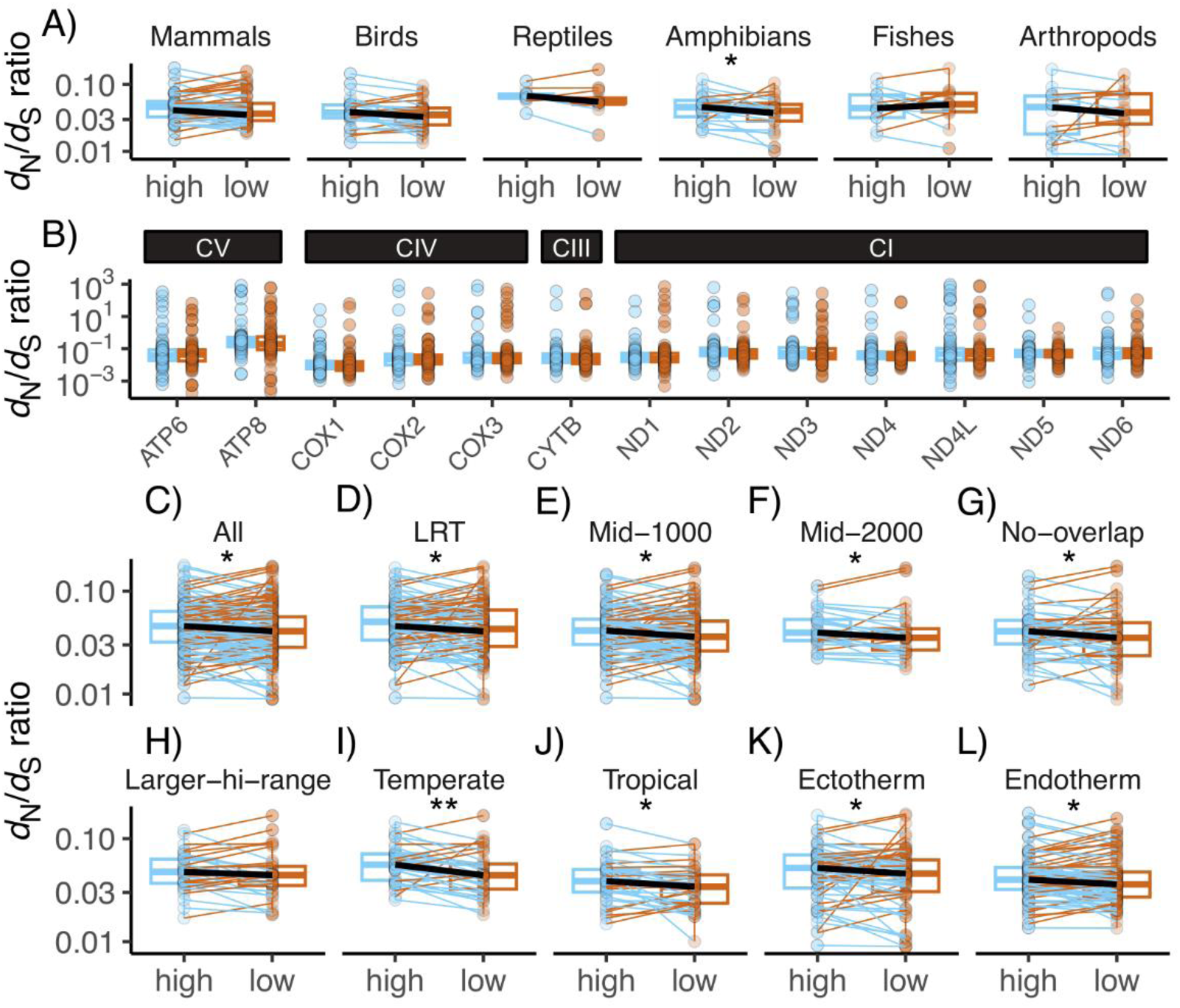
*d*N/*d*S ratios for paired high- and low-elevation species (light blue and brown, respectively) under the three-rate model. **A)** *d*N/*d*S ratios for the whole mitochondrial CDS by taxon. **B)** *d*N/*d*S ratios for all taxa by individual gene. Mitochondrial complexes to which genes belong are indicated above. **C)** *d*N/*d*S ratios for the whole mitochondrial CDS under different constraints: **All**: 154 comparisons, **LRT:** only those in which the 3-rate model is preferred to the 1-rate (null) model by likelihood ratio test, **Mid-1000:** only those in which the high and low species differ by at least 1000 m in midpoint elevation, **Mid-2000:** only those in which they differ by 2000 m, **No-overlap:** only those in which high and low species have absolutely no overlap in elevational ranges, **Larger-hi-range:** only those in which the high-elevation species has a larger range than the low, **Temperate:** only those where both taxa have midpoint latitudes above 30 °, **Tropical:** only those where both taxa have midpoint latitudes below 23.5 °, **Ectotherm:** only those among reptiles, amphibians, fishes, and arthropods, and **Endotherm:** only those among mammals and birds. Black lines connect medians while blue lines connect comparisons where the high-elevation species has higher *d*N/*d*S and brown lines connect pairs where the opposite is true. Asterisks indicate uncorrected significance by paired Wilcoxon signed-rank test (* = p < 0.05; ** = p < 0.004).

We also compared high- vs. low-elevation *d*_N_/*d*_S_ under other rate models. Values for the other models are provided in **Table S4** and supported a modestly higher *d*_N_/*d*_S_ on the high-elevation branch. However, under the two-rate model-reversed model, where the high-elevation and outgroup branches were combined, it was the low-elevation branch that had higher median *d*_N_/*d*_S_ (**Table S4**). This suggests a negative effect of overall branch length on *d*_N_/*d*_S_, with whichever taxon was combined with the outgroup having the lower *d*_N_/*d*_S_. This likely represents the well-known negative relationship between evolutionary rate and timescale, an artifact of plotting a rate against its own denominator (De Lisle & Svensson, 2023). When comparing the high- and low-elevation *d*_N_/*d*_S_ derived under the respective and opposite two-rate models, the median high-elevation rate was statistically significantly greater. This, combined with the greater difference in median *d*_N_/*d*_S_ between high- and low-taxa in the 2-rate relative to the 2-rate-reversed model, suggests a real effect of elevation on *d*_N_/*d*_S_ regardless of model selection (**Table S4**). When using AIC, the different rate models were about equally preferred: the three-rate model was preferred in 26/154 cases (16.8%), the two-rate model was preferred in 26/154 cases (16.8%), the two-rate-reversed model was preferred in 25/154 case (16.2%), the two-rate-equal model was preferred in 37/154 cases (24%), and the one-rate model was preferred in 40/154 cases (30.0%).

Median *d*_N_/*d*_S_ on the high branch under the best model by AIC for each comparison was 5.7% higher than in the low species (*P* = 0.017, *V* = 1918.5, *d.f.* = 94), supporting the existence of a high-taxon bias in *d*_N_/*d*_S_. To improve comparability across taxa and minimize branch-length artifacts, the 3-rate model was used for all subsequent comparisons.

The trend of moderately higher *d*_N_/*d*_S_ ratios in high-elevation taxa under the three-rate model held across terrestrial groups, but not in fishes (**Fig. 2A, Table S5**). Only in amphibians was the difference in medians statistically significant (*P* = 0.022, *V* = 204, *d.f.* = 22), and in no group was it significant after Bonferroni correction (*P* < 0.008). When examining differences in relative high-taxon *d*_N_/*d*_S_ between genes, there was a trend towards weakly elevated *d*_N_/*d*_S_ ratios across the mitogenome, but this was not statistically significant in any gene when analyzed by itself (**Fig. 2B, Table S6**). ATP8 tended to display a higher *d*_N_/*d*_S_ ratio than other genes regardless of elevation (**Table S6**).

We constrained our dataset of all triads to several subsets to test whether *d*_N_/*d*_S_ was consistently higher in high-elevation species (**Fig. 2C-L**). In all cases but one, median *d*_N_/*d*_S_ was statistically significantly higher among the high-elevation species prior to Bonferroni correction (**Table S7**). This included subsets where the difference in midpoint elevations was at least 1000 m (**Fig. 2E**) or 2000 m (**Fig. 2F**) or the pairs did not overlap at all in elevation (**Fig. 2G**), and subsets of only temperate taxa (**Fig. 2I**), tropical taxa (**Fig. 2J**), ectothermic taxa (**Fig. 2K**), or endothermic taxa (**Fig. 2L**). The only subset in which the difference in median *d*_N_/*d*_S_ was not statistically-significantly different was across pairs where the high-elevation species had a larger range size than its low-elevation sister species (*P* = 0.386, *V* = 316, *d.f.* = 33; **Fig. 2H**). After Bonferroni correction (*P* < 0.004), no comparison was statistically significant except for that within temperate species (*P* = 0.0036, *V* = 527, *d.f.* = 36; **Fig. 2I, Table S7**).

### Relative elevation and range size predict relative d_N_/d_S_

We used high- and low-taxon *d*_N_/*d*_S_ values from the three-rate model as the response variable in phylogenetic least-squares regressions against ecological variables. Among the high-elevation species, range-wide elevational midpoint was not a significant predictor of *d*_N_/*d*_S_ (*P* = 0.756, residual d.f. = 136, *T* = 0.3110; PGLS). However, when examining relative rather than absolute values, there was a weak but statistically significant phylogenetic correlation between the log-transformed ratio of the high/low-taxon elevational midpoints and the ratio of their *d*_N_/*d*_S_ values (PGLS; *P* = 0.021, residual d.f. = 136, *T* = 2.326; **Figure 3A**). For a 50% increase in relative elevation, the ratio between *d*_N_/*d*_S_ values increased by 1.4%. There was no significant interaction with taxon (*P* = 0.877, d.f. = 4, *F* = 0.3020; ANOVA) and including taxon as a variable did not improve the fit of the model (AIC = 309.4 without taxon, 312.2 with taxon).

**Figure 3:**
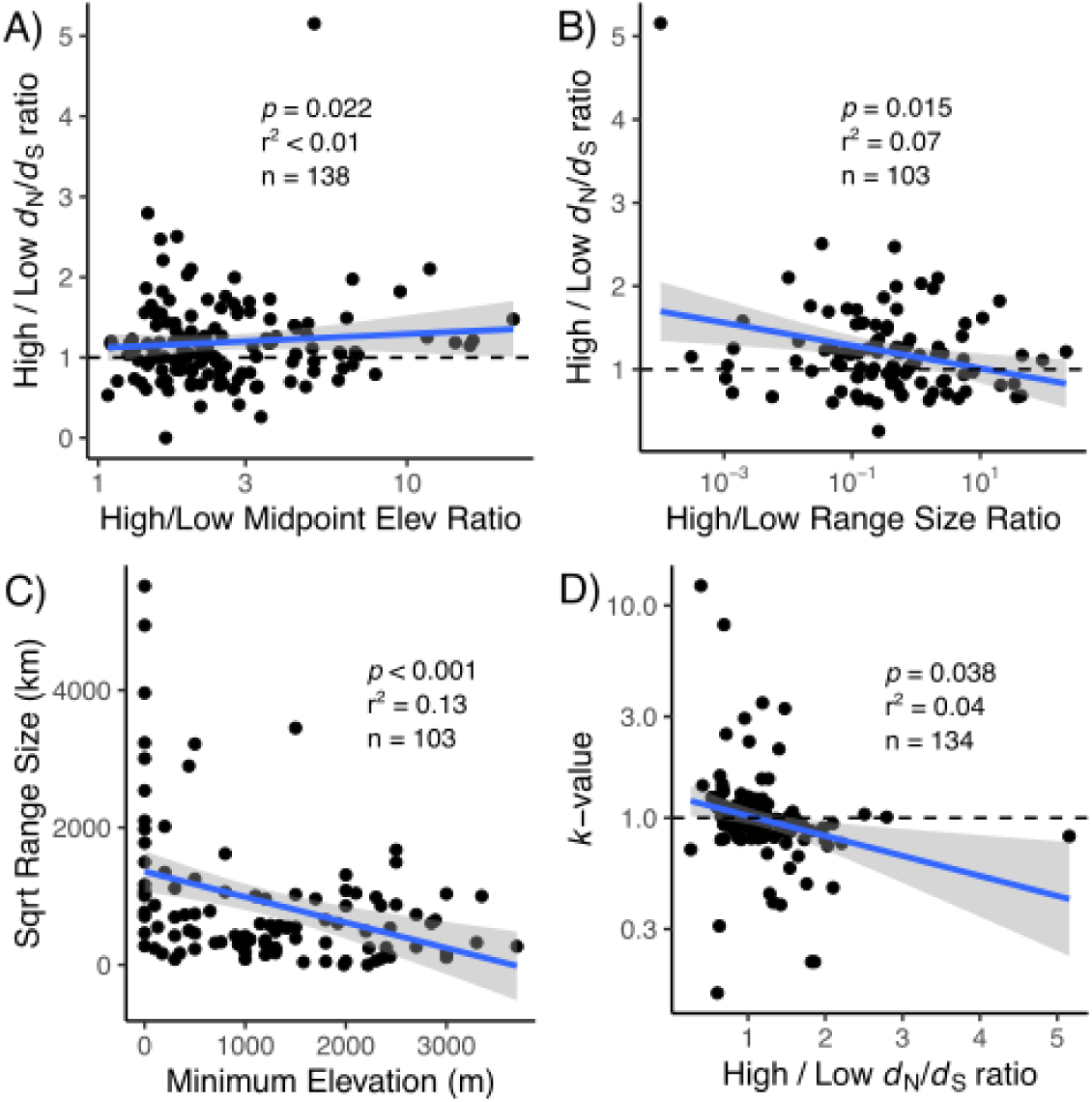
Molecular evolution in animal mtDNA as predicted by ecological correlates. **A)** Relative high-species *d*N/*d*S increases with relative high-species midpoint elevation in a phylogenetically-controlled correlation using whole-CDS *d*N/*d*S ratios. **B)** Relative high-species *d*N/*d*S decreases with relative high-species range size under a similar PGLS model. **C)** The square-root of range size decreases with minimum elevation under a linear model, illustrating how higher-living species have smaller ranges. Shading around lines of best fit show 95% confidence intervals. **D)** *k*-values decrease with relative high-species *d*N/*d*S under PGLS.

Like elevational midpoint, there was no correlation between high-taxon latitudinal midpoint and *d*_N_/*d*_S_ under the three-rate model (*P* = 0.181, residual d.f. = 105, *T* = 1.346; PGLS). There was also no correlation between the ratio of latitudinal midpoints and the ratio of *d*_N_/*d*_S_ values (*P* = 0.161, residual d.f. = 105, *T* = -1.411; PGLS). There was no correlation between log-transformed range size and *d*_N_/*d*_S_ values (*P* = 0.427, residual d.f. = 100, *T* = 0.798; PGLS). However, there was a weak but statistically significant negative correlation between the log-transformed ratio of high/low-taxon range sizes and the ratio of *d*_N_/*d*_S_ values (*P* = 0.015, residual d.f. = 100, t = -2.474; PGLS; **Figure 3B**). For a 50% decrease in relative range size, this resulted in a 0.015% increase in relative *d*_N_/*d*_S_. There was no significant interaction with taxon (ANOVA; *P* = 0.897, d.f. = 3, *F* = 0.198) and its inclusion did not improve the fit of the model (AIC = 209.7 without taxon, 217.8 with taxon). The square-root of range size decreased strongly with minimum elevation among high-elevation taxa, as predicted (*P* < 0.001, *F* = -15.31, d.f. = 100, adj. *R^2^* = 0.1241; **Figure 3C**).

### Higher relative d_N_/d_S_ values are associated with relaxed selection

There was a negative correlation between the natural log of *k*, the parameter for relaxed selection, and the high/low ratio of *d*_N_/*d*_S_ values (*P* = 0.038, residual d.f. = 131, *T* = -2.102; PGLS; **Figure 3D**), suggesting that with relatively greater high-species *d*_N_/*d*_S_, selection was more likely to be relaxed rather than intensified on high-elevation mtDNA. There was no significant interaction with taxon (*P* = 0.999, d.f. = 4, *F* = 0.028; ANOVA).

Log-transformed *k*-values were not predicted either by the log-transformed high/low ratio of midpoint latitudes (*P* = 0.841, residual d.f. = 104, *T* = -0.201; PGLS) or by the log-transformed high/low ratio of midpoint elevations (*P* = 0.541, d.f. = 134, *T* = -0.612; PGLS). Log-transformed K-values were also not predicted by the log-transformed high/low ratio of range sizes (*P* = 0.256, residual d.f. = 99, *T* = - 1.142; PGLS) or the log-transformed ratio of high/low body sizes (*P* = 0.302, residual d.f. = 115, *T* = 1.037; PGLS).

### Body size predicts absolute d_N_/d_S_ among endotherms

Because body size was measured as mass for endotherms and length for ectotherms, we did not perform a regression of absolute body size with *d*_N_/*d*_S_ from the three-rate model across the whole dataset. However, among the endotherms, the relationship between log-transformed body mass and *d*_N_/*d*_S_ was positive and highly statistically significant (*P* < 0.001 residual d.f. = 78, *T* = 4.708; PGLS; **Figure 4A**). Taxon was not a significant co-variate (*P* = 0.543, d.f. = 4, F = 0.789; ANOVA) and did not improve the fit of the model (AIC = -483.6 without taxon, -424.1 with taxon). Among the ectotherms, the relationship between body length and *d*_N_/*d*_S_ was negative and nearly statistically significant (*P* = 0.062, residual d.f. = 37, *T* = 1.921; PGLS; **Figure 4B**). We also examined how relative body size (high/low) affected relative *d*_N_/*d*_S_ values across the entire, combined dataset of endotherms and ectotherms. Relative body size across the combined dataset did not significantly predict relative *d*_N_/*d*_S_ values (*P* 0.315, residual d.f. = 117, *T*= 1.009; PGLS; Figure S2).

**Figure 4:**
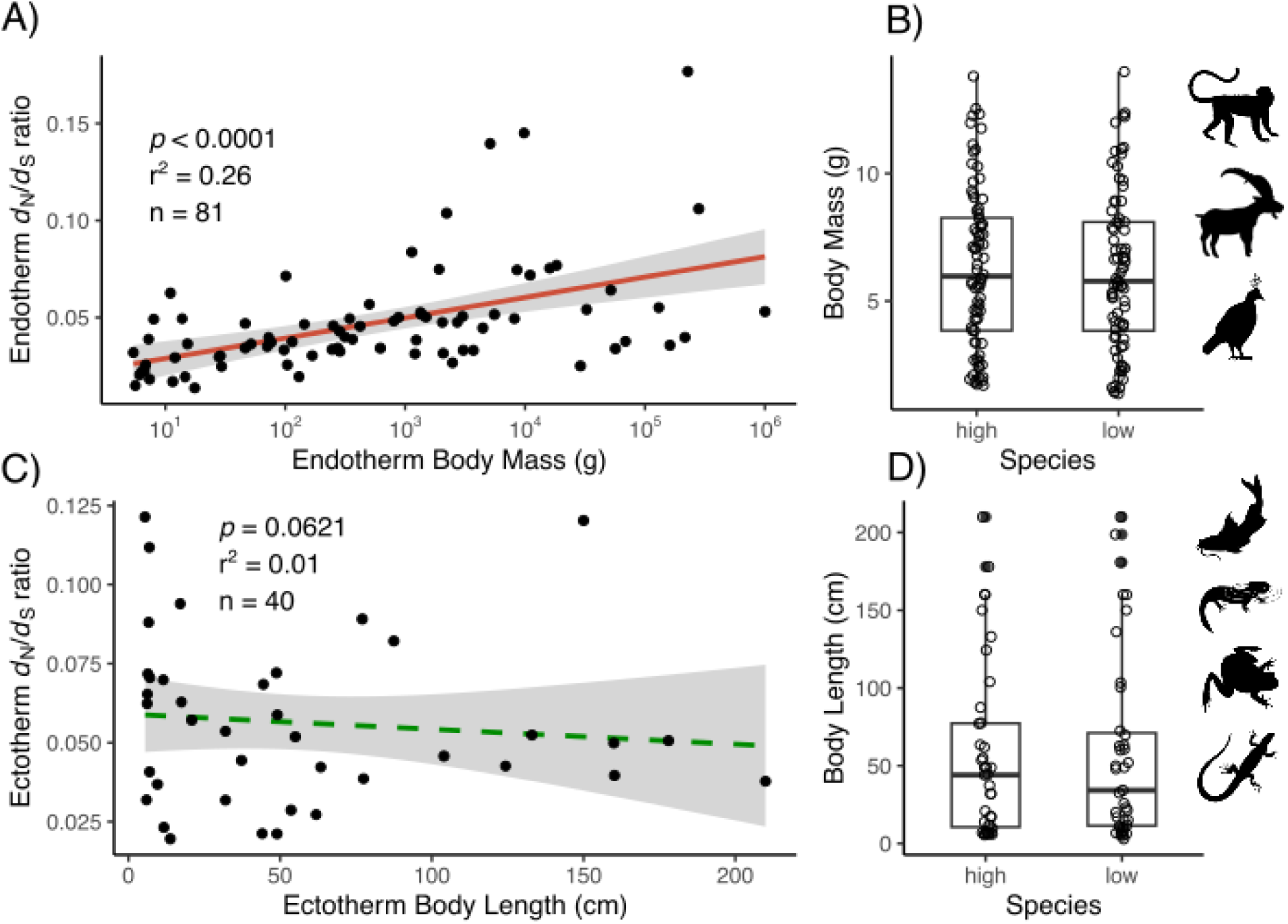
*d*N/*d*S ratios relative to body size among high-elevation **A)** endotherms and **C)** ectotherms. Endotherms have a highly significant positive relationship between log body mass and *d*N/*d*S while ectotherms have a nearly significant negative relationship between body length and *d*N/*d*S. Box plots show lack of a difference in body size between high- and low-elevation **B)** endotherms and **D)** ectotherms, suggesting trends are not a consequence of confounding elevation with body size. Silhouettes by Nat Jennings.

### Selection across clades

We also examined how selection varied with ecological factors across 6 families and suborders, finding that the relationship between ecological factors, *d*_N_/*d*_S_, and k-values varied among and within clades (**Table S8; File S4**). In nearly all analyses (23/24), multi-rate *d*_N_/*d*_S_ models were preferrable to the null, 1-rate model by likelihood ratio test (LRT) at *P* < 0.05, and after Bonferroni correction (*P* < 0.002; 22/24 cases). There was, however, a large degree of variability (*P*-values varied by over 200 orders of magnitude). Most of the variation was among the 6 taxonomic groups, with the Cercopithecidae displaying the most support for multi-rate models and Phasianoidea the least. The comparisons with the most support for the multi-rate model, such as Cercopithecidae binned by *CI* or Ranoidea by elevation, tended to have one or several highly variable bins, suggesting the highly significant likelihood ratio tests reflect the accommodation of this variability. These were not, however, necessarily the comparisons with the best evidence for linear trends between molecular evolution and ecological parameters. Significance of *k*-values is presented in terms of each bin, reflecting the results of the likelihood ratio test for relaxed selection among those taxa.

We calculated relative *d*_N_/*d*_S_ of each equal-width bin as the *d*_N_/*d*_S_ for that bin divided by the *d*_N_/*d*_S_ for the whole phylogeny under the null (one-rate) model. We visualized the trend in relative *d*_N_/*d*_S_ across bins within a clade with a linear regression weighted by the number of taxa in that bin. Because the binning scheme was not compatible with PGLS, we discuss the change in *d*_N_/*d*_S_ in terms of trends without testing for statistical significance of these linear regressions. For the same reason we also discuss trends in *k*-values across the parameter space weighted by sample size rather than statistical significance of these linear models.

Relative *d*_N_/*d*_S_ ratios tended to increase or remain constant with mid-point elevation for all groups, supporting the triad analyses suggesting moderately elevated *d*_N_/*d*_S_ ratios among high-elevation taxa (**Figure 5A**). *k*-values tended to decrease or remain constant with increasing elevation in 5/6 groups and increased in one, the Ranoidea (**Figure 5B**). Based on *k*-value likelihood ratio tests, selection was significantly intensified at low elevations in Cercopithecidae and significantly intensified at high elevations in Ranoidea; by contrast, selection was significantly relaxed at high elevation in 3/6 groups, the Bovidae + Moschidae, Cryptobranchoidea, and Lacertoidea.

**Figure 5:**
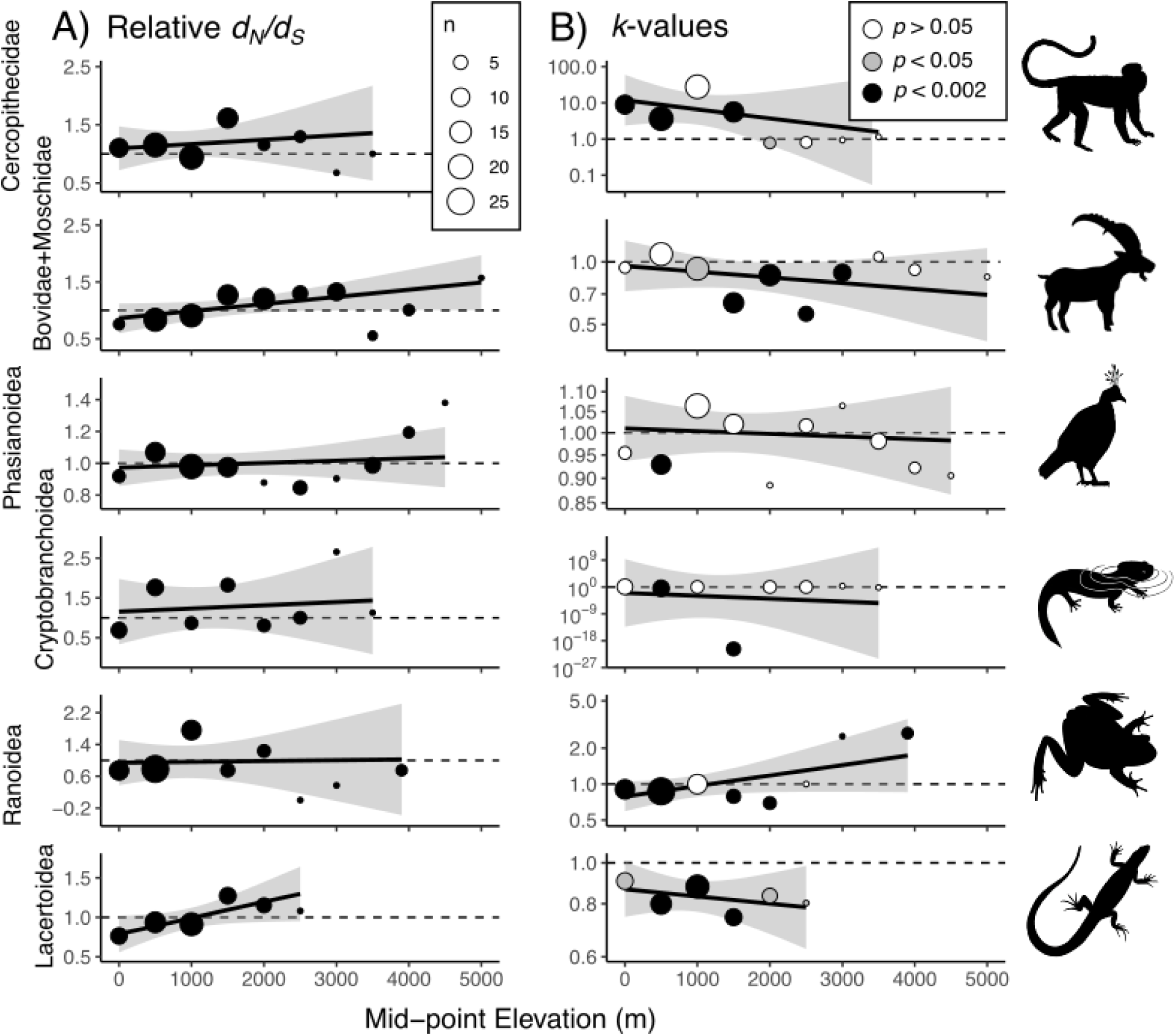
Trends in *d*N/*dS* and *k*-values across mid-point elevation in 6 family- or sub-order level clades. **A)** *d*N/*d*S ratios (y-axis) for species binned by midpoint elevation, relative to the null rate. The dashed line at y = 1 indicates equivalency between the bin rate and the null rate; points above this line are relatively higher *d*N/*d*S, indicative of intensified positive selection or relaxed purifying selection. The size of the points is proportional to the number of species in the bin. Solid lines are linear trend lines weighted by the number of species. **B)** *k*-values (y-axis) of each bin relative to the internal branches of the phylogeny. Points above the dashed line at y = 1 are indicative of intensified selection (intensified purifying and/or positive selection) while points below the line are indicative of relaxed selection. The size of points is proportional to the number of species in the bin. White points had non-significant *k*-values, while gray points were significant at *P* < 0.05 and black points were significant after Bonferroni correction (*P* < 0.002). Silhouettes by Nat Jennings.

With increasing mid-point latitude, relative *d*_N_/*d*_S_ ratios tended to increase in Bovidae + Moschidae and Lacertoidea while decreasing or remaining constant in the other groups (**Figure S1A**). *k*-values tended to increase with latitude in Cercopithecidae, Phasianoidea, and Ranoidea, but to decrease in the other three groups (**Figure S1B**). Based on *k*-value likelihood ratio tests, selection was significantly relaxed at high latitudes in Bovidae + Moschidae, Cryptobranchoidea, and Lacertoidea, while being significantly intensified at high latitudes in Phasianoidea and Ranoidea. Ranoidea also showed evidence for significantly relaxed selection at low latitudes, while in Cercopithecidae, both relaxed and intensified selection were significant at low latitudes.

Relative *d*_N_/*d*_S_ ratios tended to increase with increasing climatic index (incorporating both elevation and latitude) in Bovidae + Moschidae and Lacertoidea, while remaining relatively flat in the other groups (**Figure S2A**). *k*-values tended to increase with climatic index in Phasianoidea, Cryptobranchoidea, and Ranoidea, were relatively static in Cercopithecidae, and tended to decrease in Bovidae + Moschidae and Lacertoidea (**Figure S2B**). Based on *k*-value likelihood ratio tests, selection was significantly intensified at high climatic indexes in Ranoidea while being significantly relaxed at high climatic indexes in Bovidae + Moschidae and Lacetoidea. In Phasianoidea, selection was significantly relaxed at low climatic indexes, while in the other two groups trends were more ambiguous.

With increasing range size, relative *d*_N_/*d*_S_ ratios tended to increase in Cercopithecidae, Cryptobranchoidea, and Ranoidea while decreasing in the other three groups (**Figure S3A**). *k*-values tended to increase with range size in Cercopithecidae, Bovidae + Moschidae, Phasianoidea, and Cryptobranchoidea, while decreasing in Ranoidea and remaining relatively static in Lacertoidea (**Figure S3B**). Based on *k*-value likelihood ratio tests, selection was significantly relaxed in the smallest ranges in Bovidae + Moschidae, Phasianoidea, and Cryptobranchoidea, while in Lacertoidea selection was significantly relaxed in some bins across all range sizes. Selection was significantly intensified at the largest ranges in Cercopithecidae, Phasianoidea, and Cryptobranchoidea, while in Ranoidea responses were highly idiosyncratic: selection was significantly intensified at the smallest range sizes, and both intensified and relaxed in different bins of the largest range sizes.

## Discussion

### d_N_/d_S_ ratios tend to increase with elevation

We detected a modest but widespread signature of elevated *d*_N_/*d*_S_ ratios in the mtDNA of high-elevation species relative to their low-elevation sister taxa across several comparisons. This effect holds for almost all taxonomic groups, across mitochondrial genes, and is consistent in magnitude and direction under different inclusion criteria (**Figure 1C**). This indicates a weak but pervasive bias in nonsynonymous substitution rates in the mitochondrial CDS of high-elevation species, echoing previous results describing this pattern in specific species, within certain families, and among vertebrates as a whole (Hassanin et al., 2009; Wang et al., 2021; Xu et al., 2005) and suggests that it may be a general pattern at least for terrestrial animals.

Importantly, this modest increase in *d*_N_/*d*_S_ could reflect either positive or relaxed selection. For positive selection, we would expect *d*_N_/*d*_S_ to be more elevated in more extreme cases. While relative elevation difference weakly predicted relative *d*_N_/*d*_S_ across the triads (**Figure 3A**), constraining the dataset to stricter cases (such as large differences in midpoint elevations) barely changed median *d*_N_/*d*_S_ ratios or the difference between high and low medians (**Figure 1E-G, Table S7**). In the clade-based analysis, all groups had flat or increasing trends in *d*_N_/*d*_S_ with elevation, but only in the Ranoidea were these values associated with increasing *k*-values and significantly intensified selection (**Figure 4B**).

However, the bins that had significantly intensified selection also had *d*_N_/*d*_S_ ratios below the null model (**Figure 4A**). This indicates that intensified purifying, not positive selection had taken place in high-elevation species in Ranoidea. Purifying selection in high-elevation mitochondrial DNA is a potential signature of selection for energetic efficiency and has been documented in some groups (Graham et al., 2024).

By contrast, all other groups showed flat or decreasing trends in *k* with elevation (**Figure 4B**), suggesting increased *d*_N_/*d*_S_ ratios at high elevations may be caused more generally by relaxed, not positive selection. Selection was significantly relaxed at higher elevations in the Cercopithecidae, Bovidae + Moschidae, Cryptobranchoidea, and Lacertoidea. These significantly relaxed *k*-values on the high end of the elevation spectrum are from bins with relatively high *d*_N_/*d*_S_ ratios, providing evidence for relaxed purifying, not intensified positive selection in these groups. An association of relatively high *d*_N_/*d*_S_ values with higher *k*-values within the same bin would have been strong evidence for intensified positive selection, but was not found in any group.

Adaptation might also be expected to result in similar trends for species living at high latitudes as for those living at high elevations, if temperature is an important environmental variable in high-elevation adaptation. However, there was no effect of absolute or relative latitude on *d*_N_/*d*_S_ ratios across the triads, and relationships were mixed among the clade analyses (**Figure S3**). Among clades, no group showed unambiguous evidence for positive selection (elevated *d*_N_/*d*_S_ and significantly intensified *k*-values) with midpoint latitude or the climatic index. Responses of *d*_N_/*d*_S_ among clades to the Climatic Index, which incorporates both elevation and latitude, tended to resemble responses to midpoint elevation more than latitude (**Figure S3A,S4A**), perhaps suggesting a greater impact of environmental variation attributable solely to elevation over temperature on *d*_N_/*d*_S_ responses.

If adaptation does play a role in shaping mitogenome-wide *d*_N_/*d*_S_ ratios at high elevation, then our results suggest that oxygen limitation might be a more important selective pressure on mtDNA than temperature. In support of this, freshwater fishes are the only group of triads that trended towards higher *d*_N_/*d*_S_ ratios among low-elevation, rather than high-elevation, species. While high-elevation waters still tend to be more poorly oxygenated than those at low elevation, the increase in solubility of gasses in cold water partially compensates for the decrease in atmospheric oxygen pressure (Jacobsen & Dangles, 2017a, 2017b). Tumbling mountain rivers experienced by some high-elevation species may tend to be well-oxygenated, while lowland waters experienced by their relatives are sluggish and prone to hypoxia. Therefore, even intermittent low-oxygen conditions could be driving a signal of mt adaptation in low-rather than high-elevation fishes. In support of this, mt adaptation for greater oxygen affinity appears to play a role in adapting fishes to warmer waters (Chung et al., 2017). Unfortunately, the lack of distributional information for freshwater fish and empirical oxygen series from water at different elevations limited our ability to assess this hypothesis in a clade-based framework.

Convergent adaptation in response to elevation might be expected to result in elevated *d*_N_/*d*_S_ in one or a few specific genes across many taxa. However, relaxed selection in high-elevation species due to lower *N_e_* should be a genome-wide phenomenon affecting all genes (including nuclear-encoded genes). Our finding of similar patterns in high vs. low elevation *d*_N_/*d*_S_ values across all mt genes (**Figure 1B**) might therefore better reflect relaxed selection at high elevation. However, because the mt genome is one non-recombining linkage group, this result could also reflect genetic draft (hitchhiking) in mt genes rather than genome-wide relaxed selection (Hill, 2020). An alternative explanation is that positive selection on different genes in different lineages has resulted in similarly elevated mitogenome-wide *d*_N_/*d*_S_. Within genes there was high variability between species and *vice versa*, potentially masking important differences in the strength of selection for individual cases. Moreover, our method examining *d*_N_/*d*_S_ across whole genes and mitogenomes prevents the identification of positive selection on one or a few critical amino acid residues that may drive adaptation to high elevation. Relatively few studies have pinpointed individual amino acid substitutions in mtDNA and supported them with plausible adaptive scenarios at high elevation (see Ji et al., 2012; Kostin & Lavrenchenko, 2018; Scott et al., 2011) and adaptation may often occur via different substitutions in the genes of different species (Natarajan et al., 2016; Storz, 2017). For instance, a recent analysis showed almost no repeatability of substitutions in relation to high-elevation adaptation in the mtDNA of birds, nor in relation to several other selective pressures (Burskaia et al., 2021). The low-repeatability of molecular adaptation to the environment probably reflects the diversity of functional alterations that can be achieved within even one enzyme (Natarajan et al., 2016).

### Relaxed selection as an alternative hypothesis for elevated d_N_/d_S_ ratios

We explicitly investigated relaxed selection within triads by calculating *k* values and examining the relationship between *d*_N_/*d*_S_ and *k*. While *k* values were generally centered around 1, providing little support for an overall pattern of either relaxed or intensified selection in high-elevation species, relatively high *d*_N_/*d*_S_ values were correlated with relatively lower *k* values (**Figure 2C**). This suggests that greater relative *d*_N_/*d*_S_ ratios among high-elevation species are associated with more relaxed selection, supporting the results discussed above from our clade-based analyses.

This evidence for relaxed selection is supplemented by indirect evidence from the relationship between *d*_N_/*d*_S_ and several ecological variables which are reflective of *N_e_*. A positive relationship between *N_e_* and the effectiveness of selection is predicted from theory (Otto & Whitlock, 1997), and relaxed selection has been detected in lineages undergoing reductions in effective population size, such as arthropods transitioning to eusociality (Chak et al., 2021; Weyna & Romiguier, 2021). We found a strong, negative relationship between minimum elevation size and range size (**Figure 2D**), illustrating how species lose habitat area when constrained to higher elevations and thus may experience a reduction in *N_e_*. Perhaps as a consequence, relative range size was negatively correlated with relative *d*_N_/*d*_S_ across our triads, with high-elevation taxa that have smaller ranges showing relatively greater *d*_N_/*d*_S_ (**Figure 2B**). Additionally, the pattern of high *d*_N_/*d*_S_ values at high elevations among triads was robust to all sub-setting, except for one scenario – when the dataset was constrained to only triads where the high-elevation species had a *larger* range size, there was no longer a significant difference between high- and low-species *d*_N_/*d*_S_ (**Figure 1H, Table S7**). This illustrates that when high-elevation species experience larger range sizes than their low-elevation relatives, they do *not* display elevated *d*_N_/*d*_S_ ratios. Many of these species live on the vast Qinghai-Tibetan Plateau, where adjacent lower elevation habitats occupy smaller areas (see **Table S1**). A recent study explicitly tested for a difference in effective population size between high- and low-elevation populations of Andean ducks (Graham et al., 2024). They found no difference in *N_e_* in this guild and also no signature of positive or relaxed selection; rather, they found evidence of purifying selection in the high-elevation populations.

While relative range size did not predict *k*-values across triads, results from the clade-based analysis lent support to a relationship between the two. In the Bovidae + Moschidae, Phasianoidea, and Lacertoidea, bins with the smallest range sizes had elevated *d*_N_/*d*_S_ and significantly relaxed *k*-values (**Figure S5B**). In two other groups, Cercopithecidae and Cryptobranchoidea, bins with the largest range sizes had evidence for positive selection, part of the same hypothetical continuum. Thus, there is support from 5/6 groups for a positive relationship between range size and the effectiveness of selection. Together, these findings suggest that faster mtDNA evolution in high-elevation species may be caused more by relaxed selection than adaptive evolution.

Across our triads, endotherms exhibited a strong, positive relationship between body size and *d*_N_/*d*_S_, which may also be a signal of relaxed selection due to low effective population sizes (**Figure 3A**). Body-size has a well-supported inverse relationship with population size due to metabolic scaling relationships and habitat carrying capacities (Cotgreave, 1993; Savage et al., 2004). However, among ectotherms the relationship between body size and *d*_N_/*d*_S_ was non-significant (**Figure 3B**). In ectotherms the relationship between body size and population size is apparent at warm temperatures, but less apparent at cool temperatures where ectotherm energetic needs are lower (Buckley, Rodda, & Jetz, 2008). The metabolic flexibility of ectotherms and the intentional inclusion of many high-elevation and high-latitude species in our dataset may explain why only endotherms show the expected relationship between body size and *d*_N_/*d*_S_. Additionally, if environmental adaptation rather than low population size had been driving *d*_N_/*d*_S_ ratios among high elevation species, we may have expected smaller endotherms to exhibit the strongest signals of selection (rather than the lowest *d*_N_/*d*_S_ ratios) because of their higher mass-specific metabolic rates and surface-area to volume ratios.

Overall, we suggest that relaxation of purifying selection at high elevation is the most likely cause of modestly increased rates of overall mtDNA evolution. Small range sizes and low *N_e_* are likely when transitioning upward in elevation range, and might thus provide a better explanation for mitogenome-wide elevated *d*_N_/*d*_S_ than pervasive positive selection, which would be less efficient on all loci. However, this does not preclude the possibility that a number of important substitutions have also have been fixed by positive selection, as has been functionally demonstrated in a number of cases (Ji et al., 2012; Kostin & Lavrenchenko, 2018; Scott et al., 2011).

### The importance of distinguishing positive from relaxed selection

Tests of relaxed selection paired with *d*_N_/*d*_S_ estimates can distinguish between positive, relaxed, and purifying selection and neutrality, allowing discrimination among competing hypotheses. Without this framework, alternative stories can be told from the same data and opposing datasets can be used to support the same hypothesis (Wertheim et al., 2015; Zwonitzer & Iverson et al., 2023). For example, island species are thought to have severely reduced *N_e_* relative to their mainland ancestors, and studies have detected a strong signal of elevated *d*_N_/*d*_S_ ratios among island species relative to paired mainland taxa, arguing that this was a consequence of increased fixation of slightly deleterious substitutions due to low *N_e_* (Johnson & Seger, 2001; Woolfit & Bromham, 2005). However, another study with a similar paired design found the opposite: birds from large landmasses had higher overall rates of substitution than their island counterparts, with a trend towards higher *d*_N_/*d*_S_ (Wright et al., 2009). In this case, the authors suggested that higher *d*_N_/*d*_S_ values in mainland species resulted from more effective natural selection in large populations (intensified positive selection). This illustrates how opposite results can theoretically fit the same hypothesis, making such hypotheses hard to falsify based solely on *d*_N_/*d*_S_ ratios and similar tests. Our dataset contains examples of higher *d*_N_/*d*_S_ being associated with both smaller and larger range sizes (**Figure S5**), and RELAX analyses allowed us to distinguish among relaxed and positive selection hypotheses. For example, selection was significantly relaxed in small-ranged species of Bovidae + Moschidae, Phasianoidea, and Lacertoidea, while being significantly intensified in large-ranged species of Cercopithecidae and Cryptobranchoidea (**Figure S5**).

A bias to interpret an adaptive signal in *d*_N_/*d*_S_ ratios may be especially prevalent with animal mtDNA, which is increasingly available as whole mitochondrial genomes (Zwonitzer & Iverson et al., 2023). Because mitochondrial DNA influences animal energetics (Hill et al., 2019; Norin & Metcalfe, 2019), adaptation may be invoked to explain any signal that correlates with an interesting environmental or physiological niche. For example, after reanalyzing energetic hypotheses based on elevated *d*_N_/*d*_S_ ratios in animal mtDNA, we found that many previous cases hypothesized to demonstrate adaptive mtDNA evolution may be driven by relaxed selection instead (Zwonitzer & Iverson et al., 2023). Other recent studies have also called into question the generality and adaptive significance of mtDNA substitution bias in relation to energetics (Burskaia et al., 2021; Claramunt & Haddrath, 2023). A growing body of work takes elevated mtDNA substitution rates as a *de facto* signal of adaptation to elevation (see Introduction). However, relaxed selection on mtDNA should be explicitly tested in a phylogenetic framework, as some studies have begun to do (i.e. Gutiérrez et al., 2023).

Here, we found that while *d*_N_/*d*_S_ ratios in mtDNA do tend to be slightly elevated across high-elevation animals (**Figure 1**), there is ample variation among lineages that remains to be explained and the causes of this pattern are not straight-forward. Our work does not exclude the possibility of mtDNA adaptation to elevation or cast doubt on specific studies that have highlighted particular adaptive substitutions; however, our triad- and clade-based explicit tests tended to show more evidence for relaxed selection than adaptation shaping overall mtDNA evolution at high elevation. When combined with ecological data on species ranges, our results lend support to an alternative explanation for elevated *d*_N_/*d*_S_ ratios: high-elevation species tend to inhabit smaller ranges and likely live at lower effective population sizes, causing an uptick in slightly deleterious nonsynonymous substitutions as purifying selection is slightly relaxed. This demographic scenario explains the mitogenome-wide signature of higher *d*_N_/*d*_S_ (and predicts a similar signature in nuclear loci, a testable hypothesis), without precluding adaptation that might occur at specific loci in an idiosyncratic fashion. Overall, we caution against over-interpretation of simple molecular signatures like *d*_N_/*d*_S_ and reaffirm the need to rigorously test adaptive hypotheses and support them with thorough mechanistic investigation.

## Supporting information

Other Supplementary Tables & Figures

## Acknowledgments

We thank Allyson Gunderson, Nat Jennings, and Adam Zambie for help with data collection. We thank Mike Ryan, Misha Matz, Geoff Hill, Molly Schumer, and members of the Havird Lab for feedback. This work was funded by the National Institutes of Health (1R35GM142836) and the Stengl-Wyer Endowment at the University of Texas at Austin. This article was written without the use of artificial intelligence.

## Data Availability

The Python script used for retrieving, aligning, and concatenating mt CDSs is available at https://github.com/thekzwon/mito_accessions_to_full_alignment. The Python script used for extracting geospatial data from IUCN shapefiles is available at https://github.com/abbycriswell/spatial-data-processing.

