## Supplementary material for "Stronger evidence for relaxed selection than adaptive evolution in high-elevation animal mtDNA": Other Supplementary Tables & Figures

**Supplementary Methods**

*S1.1 Details on constructing high-low-outgroup triads*

High- and low-elevation taxa were roughly matched for latitude by binning the Earth into 5 general climatic bands: Equatorial (0˚-12˚ N or S), Dry-Tropical (12˚-23.5˚), Subtropical (23.5˚-35˚), Temperate (35˚-50˚), and Subarctic (50˚-66.5˚). In our dataset, only N hemisphere species had subarctic distributions, and no species had a primarily Arctic (above 66.5˚) distribution. For the limited number of migratory birds, comparisons were made between the breeding ranges as breeding conditions play a large role in constraining thermal niches (Socolar, Epanchin, Beissinger, & Tingley, 2017). Wherever possible, comparisons were made between species with adjacent (parapatric) distributions limited to the same climatic band; if not possible, species were compared only if their distributions overlapped in one or more climatic bands on the same continent. We did not include oceanic island species in comparisons with continental species. We avoided any comparison with other obvious ecological factors likely to be confounding between the two species’ mitochondrial evolutionary trajectories, such as a major difference in range-wide aridity (e.g., opposite sides of the Peruvian Andes), or the evolution of flightlessness in one taxon. Determination of the outgroup taxon to each pair of high/low-elevation species was based on the most recent published mitochondrial phylogeny for the group. Range maps for the four terrestrial vertebrate taxa were downloaded from the IUCN and range size and absolute latitudinal limits were extracted using a custom Python script. Body size metrics for the five vertebrate taxa, including fishes, were gathered from a variety of public databases (S1). Mass was used for birds and mammals, while snout-vent length was used for amphibians and non-avian reptiles.

*S1.2 Details on binning species based on continuous characters for the clade-based analysis*

For binning, ecological parameters were rounded (to the nearest hundred meters for elevation, to the nearest degree for latitude, to the nearest whole unit for the Climatic Index, and to the nearest 10^th^ of the natural log of range size). Tips were assigned to bins by being greater or equal than the bin minimum and less than the next bin’s minimum (i.e., greater than or equal to 2500 m midpoint elevation, rounded to the nearest hundred, but less than 3000 m), except in the case of range size, in which bins were arranged towards decreasing range size and species included were less than or equal to the bin maximum (i.e., less than or equal to *e*^6^ km^2^, in units of *e*^x.1^, but greater than *e*^5^).

*S1.3 Details on extracting data from IUCN shapefiles*

To obtain range size and latitude data for the six clades, range maps were downloaded as shapefiles from the IUCN red list (<https://iucnredlist.org>) and analyzed with a custom Python script (https://github.com/abbycriswell/spatial-data-processing) that uses the geospatial data library GeoPandas (v. 0.9.0). The program processes the spatial information contained in the many separate features comprising a species’ range shapefile, including annotations of activity (i.e. breeding, non-breeding, passage, vagrant, etc.) and status (i.e. extant, possibly extant, introduced, extinct, etc.). The program has two main defaults: a ‘current range’ that includes extant, possibly extant, and introduced breeding ranges and a ‘historical range’ that includes extant, possibly extant, and extinct breeding ranges, which we used for our analysis. The program takes the spatial union of these features of interest and provides a total range for each species in km^2^ and absolute latitudinal limits of this total range.

**Supplementary Results**

**Table S1** (*spreadsheet*): Taxa included in each triad; accession data; ecological & biogeographic data; sources.

**Table S2** (*spreadsheet*): Taxa included in the 6 clade-level comparisons; accession data; biogeographic data; sources; outgroups used for RAxML for each clade.

**Table S3** (*spreadsheet*): PAML & RELAX results for each model run, for each triad, for each gene and all concatenated genes; biogeographic data

**Table S4**: Paired Wilcoxon ranked-sign test results for median high- vs. low-taxon *d*_N_/*d*_S_ under different models. Only one rate is given for the null, which encompasses the entire tree. “Best-by-AIC” is a comparison of the rate that the high and low taxa assumed under whichever model had the lowest AIC value.

| Taxon | Median high-taxon *d*_N_/*d*_S_ | Median low-taxon *d*_N_/*d*_S_ | *P* | *V* | *d.f.* |
| --- | --- | --- | --- | --- | --- |
| *3-rate* |  |  |  |  |  |
| *2-rate* |  |  |  |  |  |
| *2-rate-reversed* |  |  |  |  |  |
| *2-rate equal* |  |  |  |  |  |
| *1-rate (null)* |  |  |  |  |  |
| *Best-by-AIC* |  |  |  |  |  |

**Table S5**: Paired Wilcoxon ranked-sign test results for taxonomic subsets of triads (Figure 1A)

| Taxon | Median high-taxon *d*_N_/*d*_S_ | Median low-taxon *d*_N_/*d*_S_ | *P* | *V* | *d.f.* |
| --- | --- | --- | --- | --- | --- |
| *Mammals* | **0.04647** | 0.03653 | 0.1163 | 913 | 54 |
| *Birds* | **0.03787** | 0.035005 | 0.0868 | 378 | 33 |
| *Reptiles* | **0.06593** | 0.05335 | 0.1082 | 64 | 12 |
| *Amphibians* | **0.04576** | 0.04039 | **0.02244** | 204 | 22 |
| *Freshwater fishes* | 0.04439 | **0.05066** | 0.7726 | 35 | 12 |
| *Arthropods* | **0.046185** | 0.03837 | 0.2019 | 85 | 15 |

**Table S6**: Paired Wilcoxon ranked-sign test results for specific genes across all triads (Figure 1B)

| Gene | Complex | Median high-taxon *d*_N_/*d*_S_ | Median low-taxon *d*_N_/*d*_S_ | *P* | *V* | *d.f.* |
| --- | --- | --- | --- | --- | --- | --- |
| *ATP6* | V | 0.03837 | **0.043115** | 0.1062 | 4375.5 | 132 |
| *ATP8* | V | **0.25281** | 0.19062 | 0.4417 | 2320 | 115 |
| *COX1* | IV | **0.00951** | 0.00874 | .208 | 2154 | 110 |
| *COX2* | IV | **0.023355** | 0.021595 | .2909 | 1909 | 109 |
| *COX3* | IV | 0.0229 | **0.02442** | .2191 | 3372 | 127 |
| *CYTB* | III | **0.02806** | 0.02335 | .3317 | 4513 | 138 |
| *ND1* | I | **0.02789** | 0.024945 | .2277 | 3852 | 130 |
| *ND2* | I | **0.06014** | 0.0521 | .05603 | 5314 | 139 |
| *ND3* | I | **0.051705** | 0.042885 | .2794 | 2099 | 107 |
| *ND4* | I | **0.03835** | 0.03749 | .2401 | 4912 | 140 |
| *ND4L* | I | **0.04748** | 0.04482 | .1256 | 1519 | 92 |
| *ND5* | I | **0.04924** | 0.04846 | .6046 | 4808 | 144 |
| *ND6* | I | 0.045725 | **0.053995** | .6511 | 2661.5 | 117 |

**Table S7**: Paired Wilcoxon ranked-sign test results for other subsets of triads (Figure 1C)

| Test | Description | *P* | *V* | *d.f.* |
| --- | --- | --- | --- | --- |
| *All* | All pairs | **0.006383** | 7348.5 | 153 |
| *LRT* | Pairs where the 3-rate modeled was preferred by  likelihood ratio test | **0.01943** | 2837 | 94 |
| *Mid-1000* | Pairs differing by at least 1000 m in midpoint elevation | **0.006588** | 3432 | 102 |
| *Mid-2000* | Pairs differing by at least 2000 m in midpoint elevation | **0.01585** | 297 | 27 |
| *No-overlap* | Pairs with a high-species minimum equal to or greater than the low-species maximum elevation | **0.0285** | 744 | 46 |
| *Larger-hi-*  *range* | Pairs where the high-elevation species has a larger range  size | 0.386 | 316 | 33 |
| *Temperate* | Pairs where both species have midpoint latitudes at or  above 30˚ | **0.003603** | 527 | 36 |
| *Tropical* | Pairs where both species have midpoint latitudes at or  below 30˚ | **0.01965** | 464 | 35 |
| *Ectotherm* | All pairs of mammals and birds | **0.04108** | 1339 | 64 |
| *Endotherm* | All pairs of reptiles, amphibians, arthropods, and  freshwater fishes | **0.04032** | 2430 | 88 |

**Table S8** (*spreadsheet*): PAML & RELAX results for clade-level comparisons; biogeographic data

**Supplementary Figures**

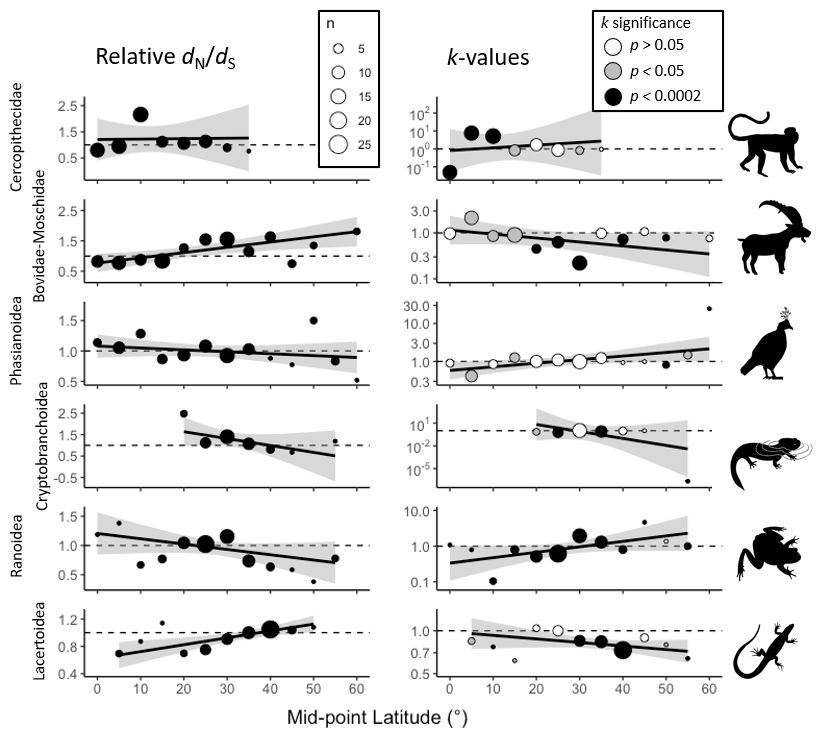

**Figure S1:** Relative *d*_N_/*d*_S_ and *k*-values, as in Figure 4, but for bins based on midpoint latitude.

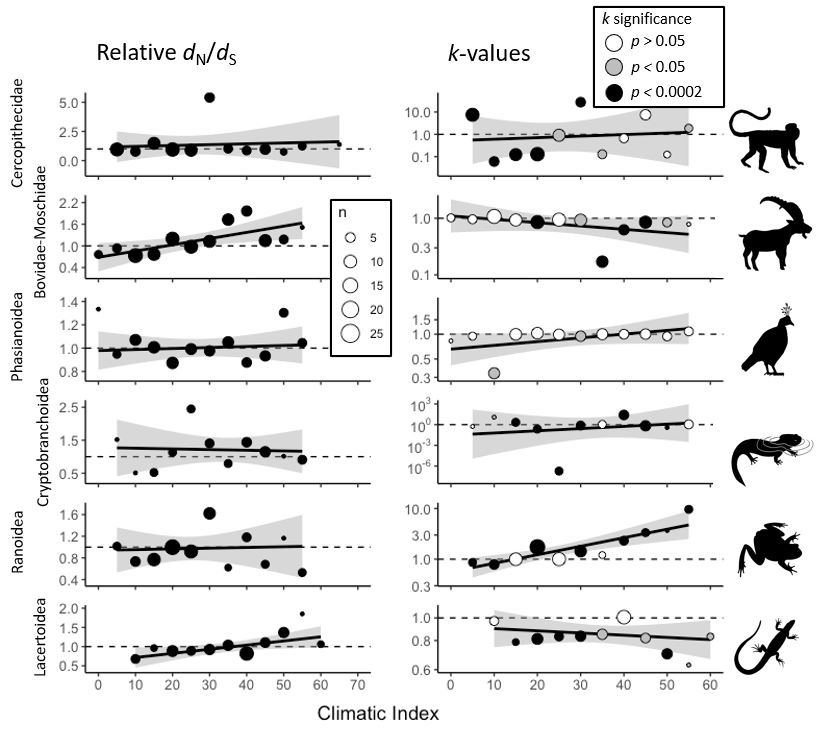

**Figure S2:** Relative *d*_N_/*d*_S_ and *k*-values, as in Figure 4, but for bins based on Climatic Index.

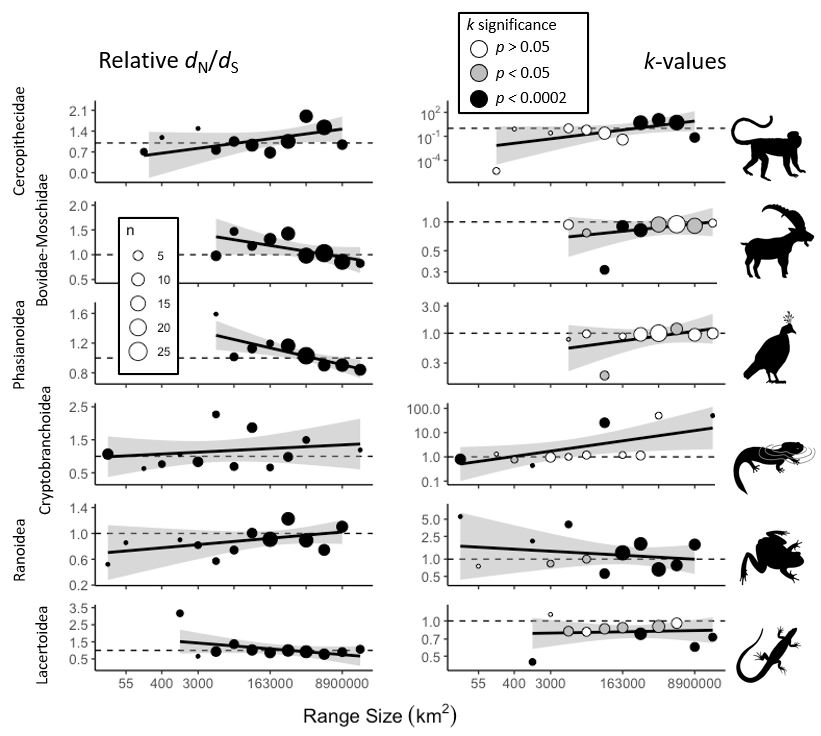

**Figure S3:** Relative *d*_N_/*d*_S_ and *k*-values, as in Figure 4, but for bins based on range size.

**Supplementary Files (pending on FigShare)**

**Supplementary File S1:** Alignments used for all analyses

**Supplementary File S2**: PAML control files & result files

**Supplementary File S3**: RAxML outputs & trees

**Supplementary File S4**: RELAX outputs
